# Shedding light on the virome of *Caliothrips phaseoli*, an emergent pest in peanut

**DOI:** 10.64898/2026.10.09.758028

**Authors:** Nicolás Bejerman, Soledad de Breuil, Carolina Dottori, Humberto Debat

## Abstract

The bean thrips, *Caliothrips phaseoli*, is an emergent pest affecting peanut production. Thus, gaining a deeper understanding of the naturally occurring viral populations associated with this insect is of major importance, as such knowledge could support the design of novel pest management strategies that are both ecologically responsible and sustainable over the long term. In this study, we performed the first RNA high-throughput sequencing (HTS) analysis of *C. phaseoli* to unravel its virome, discovering a wide number of novel RNA viruses. Our HTS analysis revealed the presence of 31 viral segments, corresponding to 18 viruses, including double-stranded RNA viruses as well as positive- and negative-sense single-stranded RNA viruses. All these are novel viruses and cover multiple viral lineages belonging to diverse families. Overall, these results indicates that *C. phaseoli* hides a complex virome. The findings presented here provide an important framework for subsequent studies aimed at understanding the ecological significance, evolutionary patterns, and possible use of these viruses as components of biologically based management programs for this emerging peanut pest.

## 1. Introduction

Thrips (Thysanoptera) are widespread agricultural pests that can cause substantial damage to crops through their feeding and oviposition activities. By disrupting plant tissues and removing cellular contents, thrips feeding can produce characteristic symptoms such as discoloration, silvering or bronzing, leaf deformation, and, under severe infestations, reduced plant growth and crop performance (Childers, 1997). Feeding and oviposition injuries may also facilitate the entry of plant pathogens into damaged tissues, while thrips can contribute to pathogen dispersal by carrying microorganisms or fungal spores on their bodies. In addition, some thrips species act as vectors of economically important plant viruses (Childers, 1997; Rotenberg et al., 2015).

Among agriculturally important thrips, the bean thrips, *Caliothrips phaseoli* (Hood) is a polyphagous species widely distributed throughout the Americas, with records from North, Central, and South America and the Caribbean (Mound and Marullo, 1996; Viteri et al., 2009; Santos and Lima, 2024). It has been reported in association with plants from several botanical families and with major crops including common bean, soybean, cotton, and peanut (Mound and Marullo, 1996; Sosa et al., 2017; Santos and Lima, 2024). Its economic importance has been particularly recognized in legume cropping systems, including common bean and soybean, in different regions of the Americas (Viteri et al., 2009; Gerardo et al., 2021; Santos and Lima, 2024).

Although *C. phaseoli* has long been recognized as an agricultural pest, recent reports indicate renewed attention to its occurrence and impact in several cropping systems. In Mexico, it is currently regarded as one of the major thrips pests of common bean, with increasing pest pressure reported particularly in important production areas (Miranda Valdez et al., 2026). In Brazil, where *C. phaseoli* is considered an important pest of common bean and soybean (Santos and Lima, 2024), high thrips infestations have been reported in soybean in the southern region since 2017, with *C. phaseoli* among the main species present (Warpechowski et al., 2024). Recent reports have also documented feeding damage by *C. phaseoli* on *Arachis* spp. (Santos and Lima, 2024) and damaging infestations on forage grasses (Torres et al., 2024).

In Argentina, *C. phaseoli* is an important component of thrips communities associated with both soybean and peanut in Córdoba Province. It is the dominant thrips species in soybean in the south-central region of the province, where feeding damage can negatively affect crop yield (Gerardo et al., 2021). In peanut, *C. phaseoli* has been documented for at least two decades, causing characteristic foliar damage (Boito et al., 2006), and is among the most abundant thrips species found in commercial crops (de Breuil et al., 2021). More recently, *C. phaseoli* has been described as the thrips pest most commonly observed in peanut in central-southern Córdoba, and a diagrammatic scale was specifically developed to quantify the foliar damage it causes to this crop (Crenna et al., 2023). Together, its broad geographic distribution, association with diverse crop hosts, and recent reports of damaging populations across distinct agricultural systems highlight the increasing attention being given to *C. phaseoli* as an agricultural pest. Despite increasing knowledge of the biology, ecology, population dynamics, and management of *C. phaseoli* (Sosa et al., 2017; Gerardo et al., 2021), the viruses associated with this species remain largely unexplored. The absence information about its virome, limits our ability to fully assess how viruses may affect host population dynamics and influence adaptation to changing environmental conditions. Viral pathogens are known to alter numerous aspects of arthropod biology, including physiology, behavior, and responses to environmental stress (Shikano et al., 2017); as a result, interactions between hosts and their associated viruses could play a major role in determining outbreak potential and may also affect the success of pest control approaches (Trebicki et al., 2017). Therefore, investigating the diversity and composition of the virome linked to *C. phaseoli* represents a critical step toward improving our understanding of this pest and supporting the development of more efficient management strategies.

The introduction of high-throughput sequencing (HTS) technologies has dramatically changed the field of virus discovery by enabling comprehensive exploration of the previously unrecognized diversity that composes the global virosphere (Kristensen et al., 2010). In contrast to conventional approaches, such as virus isolation or PCR-based detection, which rely on pre-existing sequence information, HTS permits unbiased identification of viral genomes without prior assumptions (Liu et al., 2011). This methodological advance has had a profound impact on invertebrate virology, where large-scale meta-transcriptomic analyses have revealed an enormous number of previously undescribed viruses and substantially expanded current knowledge of RNA virus evolution and phylogenetic diversity (Shi et al., 2016). More recent investigations have broadened these discoveries to marine ecosystems, uncovering an extensive and largely overlooked diversity of RNA viruses associated with aquatic invertebrates (Zhang et al., 2021). In addition, the high sensitivity and depth of HTS have facilitated detailed characterization of species-specific viromes, including the identification of complex viral assemblages in the golden orb-weaver spider, *Nephila clavipes* (Debat, 2017). Together, these advances have significantly altered current perspectives on the diversity, evolution, and ecological distribution of viruses infecting invertebrate hosts.

In the present study, we used HTS to characterize, for the first time, the virome associated with *C. phaseoli* collected from peanut crops in Córdoba, Argentina. The main objective was to identify and characterize previously unknown viruses linked to this thrips, thereby revealing an unexplored biological component that could influence multiple aspects of the insect’s ecology and population dynamics. By examining the viruses associated with this species, we aimed to establish a foundational resource for future studies focused on the ecological significance and evolutionary relationships of insect-associated viruses in agricultural environments. Expanding knowledge of the virome of *C. phaseoli* also contributes to a broader understanding of host-virus interactions in agroecosystems and may provide valuable insights into mechanisms shaping insect adaptation and pest emergence. In addition, identifying naturally occurring viruses in this species could open new opportunities for the development of biologically based and environmentally sustainable pest management approaches. Altogether, this research addresses a poorly explored aspect of a pest of major economic relevance to peanut production and represents a significant advance toward understanding its associated viral diversity. Notably, we identified 18 novel RNA viruses, which constitute, to the best of our knowledge, the first viral records described for *C. phaseoli*, highlighting the remarkable complexity and diversity of its virome.

## 2. Material and methods

### 2.1. Thrips sampling and sample processing

Four commercial peanut fields distributed throughout the central and southern areas of Córdoba Province, Argentina, were evaluated (Table S1). In each field, thrips were collected using a systematic sampling scheme. Five sampling stations were established along a diagonal transect, with stations separated by 10 crop rows. At each station, one beat per plant was performed on 10 consecutive plants within the same row, and insects were collected using a plastic tray containing 70% ethanol. In the laboratory, adult *C. phaseoli* specimens were separated and identified under a stereomicroscope based on diagnostic external morphological characters described for the species (Boito et al., 2006; Lima et al., 2020). All *C. phaseoli* specimens collected from the same field were then pooled into a single sample for nucleic acid extraction.

### 2.2. RNA Extraction and HTS Analysis

Total RNA was extracted from pools of 25 *C. phaseoli* specimens from each one the four evaluated fields. Insects were ground using a mortar and pestle in liquid nitrogen, and total RNA was extracted using Trizol (Invitrogen, USA) following manufacturer’s instructions. Resulting RNAs were treated with DNAse and sent to Macrogen INC (South Korea) for library preparation and HTS sequencing. Library was carried out using the TruSeq Stranded Total RNA Library Prep Kit with Ribo-Zero Gold (Illumina, USA). Ribodepleted RNAs were then sequenced in paired-end 101 bp read format on an Illumina NovaSeq6000 platform.

The raw paired-end sequencing reads were first processed to remove low-quality regions and adapter sequences using *Trimmomatic* (v0.40; http://www.usadellab.org/cms/?page=trimmomatic). Default settings were applied with one key modification: the quality threshold was increased from 20 to 30, including adjustments during the initial ILLUMINACLIP step and the sliding-window trimming procedure, ensuring a minimum average quality score of 30. After filtering, the high-quality paired reads were assembled *de novo* with *rnaSPAdes* under standard conditions using the Galaxy platform (https://usegalaxy.org/). The assembled contigs were subsequently compared against a comprehensive non-redundant (NR) database of viral proteins obtained from NCBI (https://www.ncbi.nlm.nih.gov/protein/?term=txid10239[Organism:exp]) using local BLASTX searches with an E-value cutoff of 1 × 10^-^□. All candidate viral hits identified in this step were carefully evaluated.

Contigs showing similarity to viral sequences were then subjected to further refinement. This included manual curation, extension, and validation through an iterative read-mapping approach. In this process, reads corresponding to each candidate contig were selectively extracted and used to extend the sequence; the updated contig was then re-used as a query in successive rounds until no further extension was possible. Finally, the resulting extended and curated transcripts were re-assembled using the alignment tool in *Geneious* (v8.1.9; Biomatters Ltd., Boston, MA, USA), applying high-sensitivity settings to ensure optimal sequence accuracy.

### 2.3 Sequence and Phylogenetic Analysis

ORFfinder with a minimal ORF length of 150 nt and genetic code 1 (https://www.ncbi.nlm.nih.gov/orffinder/) was used to predict the open reading frames (ORFs). InterPro https://www.ebi.ac.uk/interpro/search/sequence-search) and the NCBI Conserved domain database-CDD v3.20 (https://www.ncbi.nlm.nih.gov/Structure/cdd/wrpsb.cgi) with an e-value threshold of 0.01 were employed to identify the functional domains and architecture of translated gene products. In addition, HHPred and HHBlits, as implemented in https://toolkit.tuebingen.mpg.de/#/tools/, were used to complement the annotation of divergent predicted proteins using hidden Markov models. To determine the taxonomic position of each identified virus, relevant proteins were chosen according to the related viral family of each virus identified in our study. Phylogenetic analysis were carried out using MAFFT 7.526 (https://mafft.cbrc.jp/alignment/software/) with multiple amino acid (aa) sequence alignments using the best-fit model. The aligned aa sequences were used as input to generate phylogenetic trees through the maximum-likelihood method with the FastTree 2.1.11 tool available at http://www.microbesonline.org/fasttree/. Local support values were calculated with the Shimodaira-Hasegawa test (SH) and 1000 tree resamples. The sequences of proteins of relevant viral families related to those identified viruses were used as outgroup in the phylogenetic trees.

### 2.4 Metagenomic taxonomic profiling

Quality-trimmed reads from a whole-insect RNA-seq library of *Caliothrips phaseoli* were analyzed with the CZ ID (formerly IDseq) metagenomic next-generation sequencing pipeline (Kalantar et al., 2020), which performs host and quality filtering, short-read alignment to the NCBI nucleotide (NT) and non-redundant protein (NR) databases, *de novo* assembly of non-host reads, and re-alignment of contigs to assign reads and contigs to taxa. Because *C. phaseoli* genome is not available among the pipeline’s host-subtraction options, host filtering could not remove the insect’s own transcripts; this is a defining feature of the dataset. Abundances were calculated as reads per million (rPM) as defined by the pipeline, retaining for every taxon the separate NT and NR statistics (rPM, read count, assembled-contig count, mean percent identity, alignment length and E-value).

### 2.5 Eukaryotic and bacterial annotation

Eukaryotic taxa were assigned an ecological role from their lineage: thrips host (order Thysanoptera), other arthropod (phylum Arthropoda, non-Thysanoptera), plants (Viridiplantae), fungi (including entomopathogens), or other eukaryote. Bacterial taxa were assigned to a transparent confidence class to separate genuine insect associates from artefacts: “recognized insect endosymbiont” (genera *Rickettsia*, *Cardinium*, *Wolbachia*, *Arsenophonus*, *Spiroplasma*, *Hamiltonella*, *Sodalis*, *Portiera*, *Sulcia*/*Karelsulcia*, *Buchnera*, *Lariskella*, *Amoebophilus* and relatives), “likely enteric/clinical contaminant” (human- and laboratory-associated enterobacteria and clinical taxa such as *Salmonella*, *Escherichia*, *Klebsiella*, *Streptococcus*, *Staphylococcus*, *Bacteroides fragilis*), and “other/environmental”. This classification is taxonomic and is intended to make the alternative hypotheses for each bacterial detection explicit rather than to assert symbiosis.

## Results and Discussion

### 3.1 Metagenomic taxonomic profiling

Raw data analysis resulted into 8,581 classified species-level taxa: 4,455 eukaryotic, 772 bacterial, 81 viral and 19 archaeal. By nucleotide reads the sample is overwhelmingly eukaryotic (99.96% of NT rPM), with bacteria contributing 0.04% and viruses a negligible nucleotide fraction (Supp. Fig. X (T7)). Although viruses contribute almost no nucleotide reads, they sum to ∼1,600 rPM of *protein* (NR) evidence, and that evidence points to a markedly divergent virome

### 3.2. Viruses associated with C. phaseoli

Insects are the most abundant and widely distributed animal group on Earth (Stork, 2018). Advances in HTS technologies have demonstrated that these organisms host extraordinarily diverse and complex viral communities (Debat et al., 2024; Kafer et al., 2019; Li et al., 2015; Li et al., 2026; Qi et al., 2023; Shi et al., 2016; Wu et al., 2020). Nevertheless, even with the rapid expansion of sequencing-based studies, only a limited portion of the true viral diversity associated with insects has been characterized to date, indicating that the vast majority of insect viromes remains unexplored (Xavier et al., 2021; Harvey and Holmes, 2022). The viromes of just five thrips species (*Frankiniella intosa, Frankiniella occidentalis, Neohydatothrips variabilis, Stenchaetothrips biformis, Thrips palmi*) were explored so far and more than one hundred viruses were discovered (Thekke-Veetil et al., 2020; Chiapello et al., 2021; Zhang et al., 2025; Ma et al., 2026); emphasizing the potential richness of the virome associated to thrips.

More than 6500 thrips species have been identified worldwide (ThripsWiki, 2025), and these small insects display considerable variation in their biology, ecology, and hosts (Mound, 2005). Despite this diversity, current knowledge regarding the viruses associated with psyllids remains extremely limited. Available studies likely represent only a small fraction of the actual viral diversity present within this insect group, providing a fragmented and potentially biased understanding of their viromes. Consequently, it is reasonable to assume that the viral communities associated with economically significant thrips pests, including the bean thrips, are still poorly explored and largely uncharacterized. Therefore, in this study, we explored the virome of *C. phaseoli*, which led to the discovery of 18 RNA viruses (Table 1).

**Table 1.**
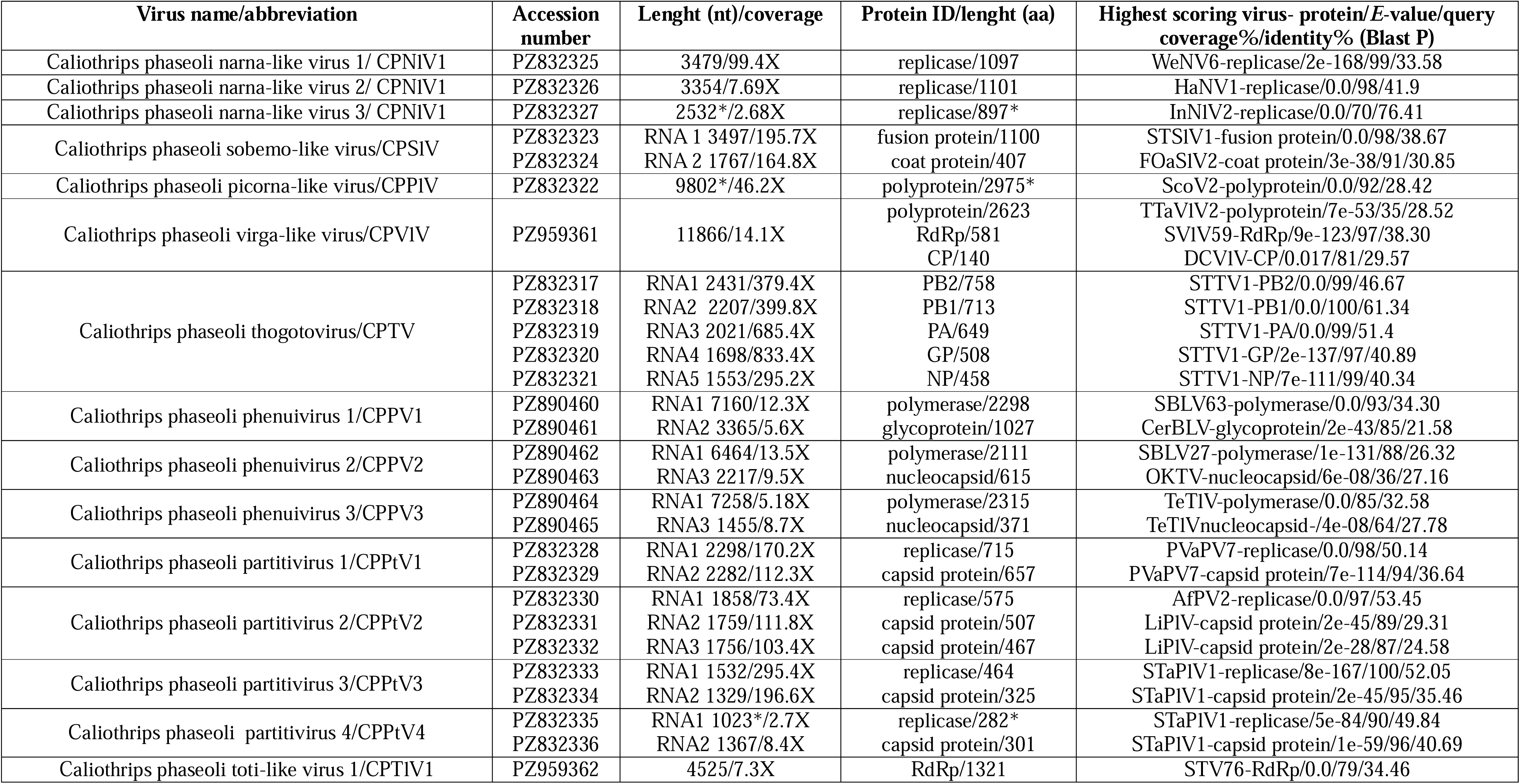
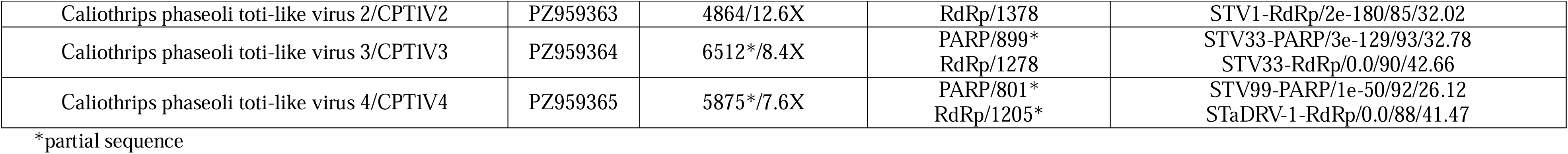
Summary of the viruses identified in *Caliothrips phaseoli*.

Our investigation of the viral community associated with *C. phaseoli* revealed a remarkably broad and complex virome linked to this economically significant pest of peanut. Expanding knowledge of the viruses infecting this pest represents an important foundation for future studies aimed at developing highly targeted biological control approaches that exploit species-specific viral pathogens to suppress thrips populations.

### 3.3. (+)ssRNA viruses

#### 3.3.1. Molecular and phylogenetic characterization of the newly identified bean thrips narna-like viruses

Three contigs shows similarity to narna-like virus sequences. Members of the family *Narnaviridae* are non-encapsidated characterized by positive-sense single-stranded RNA genomes that typically encode only one open reading frame (ORF) (Hillman and Cai, 2013). This viral family is assigned to the order *Wolframvirales*. According to the current classification established by the International Committee on Taxonomy of Viruses (ICTV), just two species are formally included in the genus *Narnavirus* (https://ictv.global/taxonomy). As a consequence, the vast majority of narna-like viruses identified so far; which are well over one thousand, have not yet been assigned to any formal taxonomic classification. Despite most of the narnaviruses reported to date are associated to fungal hosts (Sadiq et al., 2022), several of them were also discovered in insect metagenomes (Shi et al., 2016; De Risi et al., 2019; Wu et al., 2020; Chiapello et al., 2021; Qi et al., 2023; Cao et al., 2026); including some that were identified in thrips samples (Thekke-Veetil et al., 2020; Chiapello et al., 2021).The three identified narna-like viruses were named as Caliothrips phaseoli narna-like virus 1-3 (CPNLV1-3) (Table 1). The two viruses with fully assembled coding-sequences (CPNLV1 and CPNLV2) contain a single ORF encoding an RdRp (Fig. 1A), with their genome lengths composed of 3479 nt and 3354 nt, respectively (Table 1).

**Figure 1.**
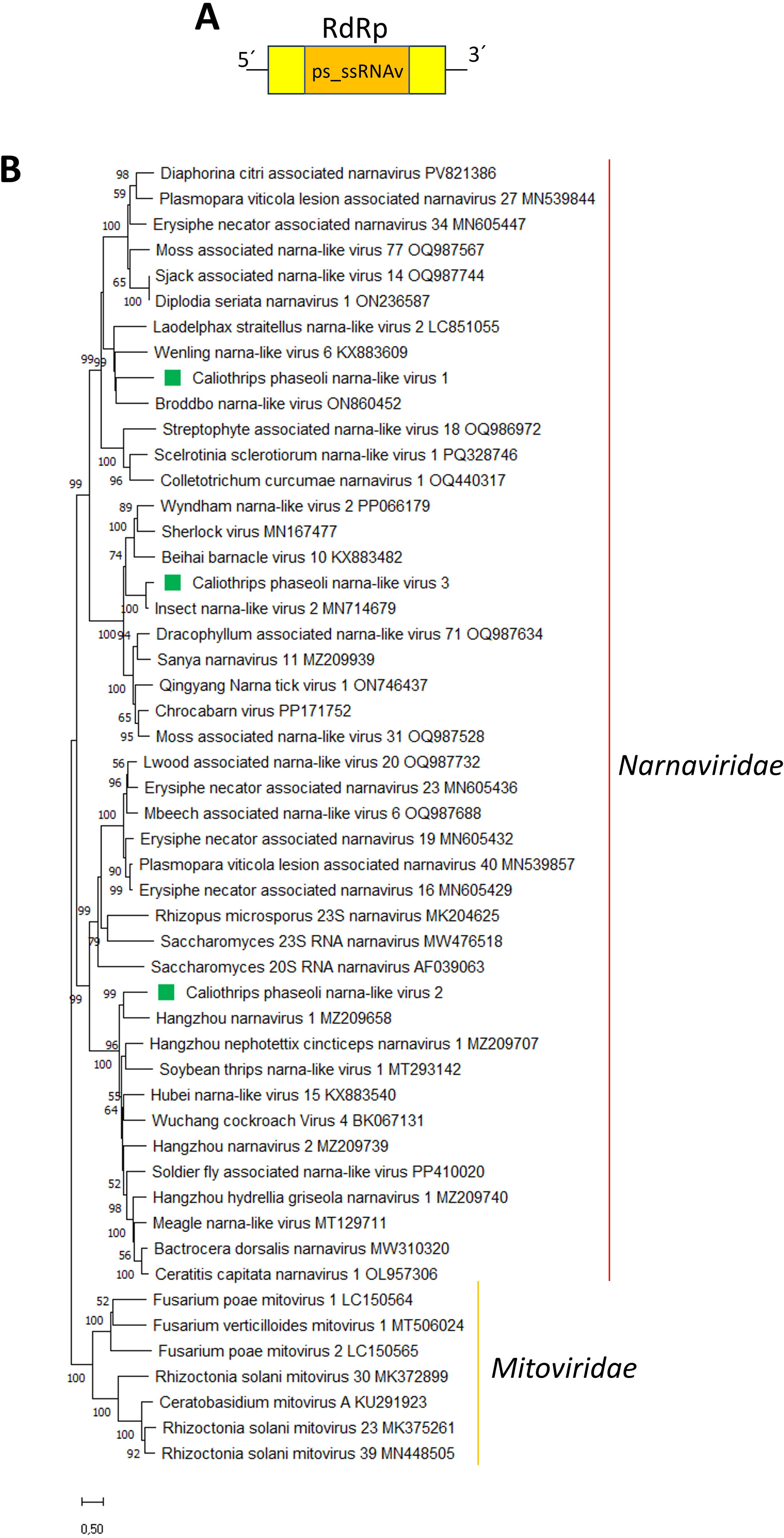
(A) Schematic representation of the genome organization of Caliothrips phaseoli narna-like virus 1-3. (B) Maximum likelihood phylogenetic trees reconstructed using the RdRp sequence of Caliothrips phaseoli narna-like virus 1-3 and of representative narna-like viruses. Bootstrap values above 50% are shown (1000 replicates). Caliothrips phaseoli narna-like virus 1-3 are indicated with a green square. The scale bar shows the substitution per site.

BLASTp searches using the deduced aa sequences showed that the best hits for CPNLV1-3 were other narna-like viruses associated to insects, with identity values below 45% for CPNLV1-2 (Table 1), and below 80% for CPNLV3 (Table 1). Phylogenetic analysis based on RdRp aa sequence alignment revealed that the three viruses were located in different clades, grouped with distinct narna-like viruses associated to insects (Fig. 1B). CPNLV1 clustered together with Broddbo narne-like virus, which was identified in mites samples (Ortiz-Baez et al., 2022), while CPNLV2 clustered together with Hangzhou narnavirus 1, which was identified in invertebrate samples collected in Hangzhou, China (Wang et al, 2024); on the other hand, CPNLV3 clustered together with insect narna-like virus 2, which as identified in invertebrate samples from unconfirmed origin (Chiapello et al., 2021) (Fig 1B). The ps_ssRNAv_RdRp conserved domain was identified in the protein encoded by CPNLV1-3 (Fig. 1A), including the GDD motif, which is present in the RdRp encoded by narnaviruses (Chiapello et al., 2021).

#### 3.3.2. Molecular and phylogenetic characterization of the newly identified bean thrips Sobemo-like virus

Two contigs show similarity to sobemo-like virus sequences. The family *Solemoviridae* was recently created, and comprises positive-sense single-stranded RNA viruses. To date, every identified member of this viral family has been associated with plant hosts (Somera et al., 2021). Members of this viral family are typically characterized by a monopartite, positive-sense RNA genome containing approximately four to ten open reading frames (ORFs). These coding regions are responsible for producing several essential viral proteins, such as a suppressor of host RNA silencing pathways, a capsid protein, and a polyprotein that harbor conserved domains, including serine protease and RNA-dependent RNA polymerase (Somera et al., 2021). Nevertheless, recent research have identified a number of sobemo-like viruses in arthropods that deviate from this canonical genomic arrangement by possessing bipartite genomes. Such viruses have been detected across a broad range of invertebrate taxa, including crustaceans (Chang et al., 2023), myriapods (Shi et al., 2016), and diverse insect species (Otatti et al., 2020; Aragao et al., 2023; Dahan et al., 2024; Debat et al., 2025; Serra et al., 2026; Candia et al., 2026).

In this study, one sobemo -like virus was identified and named as Caliothrips phaseoli sobemo-like virus (CPSlV) (Table 1), which add a novel member to the group of sobemo-like viruses with bi-segmented genomes associated to non-plant hosts. Sobemo-like viruses were also identified in the thrips species *Neohydatothrips variabilis*, *Frankliniella occidentalis* and *Thrips tabaci* (Thekke-Veetil et al., 2020; Chiapello et al., 2021).. Therefore, *C. phaseoli* is the fourth thrips host where a sobemo-like virus was identified.

The CPSlV genome is organized into two distinct segments of single-stranded RNA with positive polarity (Figure 2A and Table 1). The length of the first RNA segment is 3497 nt; while the second segment is shorter, with a lengths of 1767 nt (Table 1).

**Figure 2.**
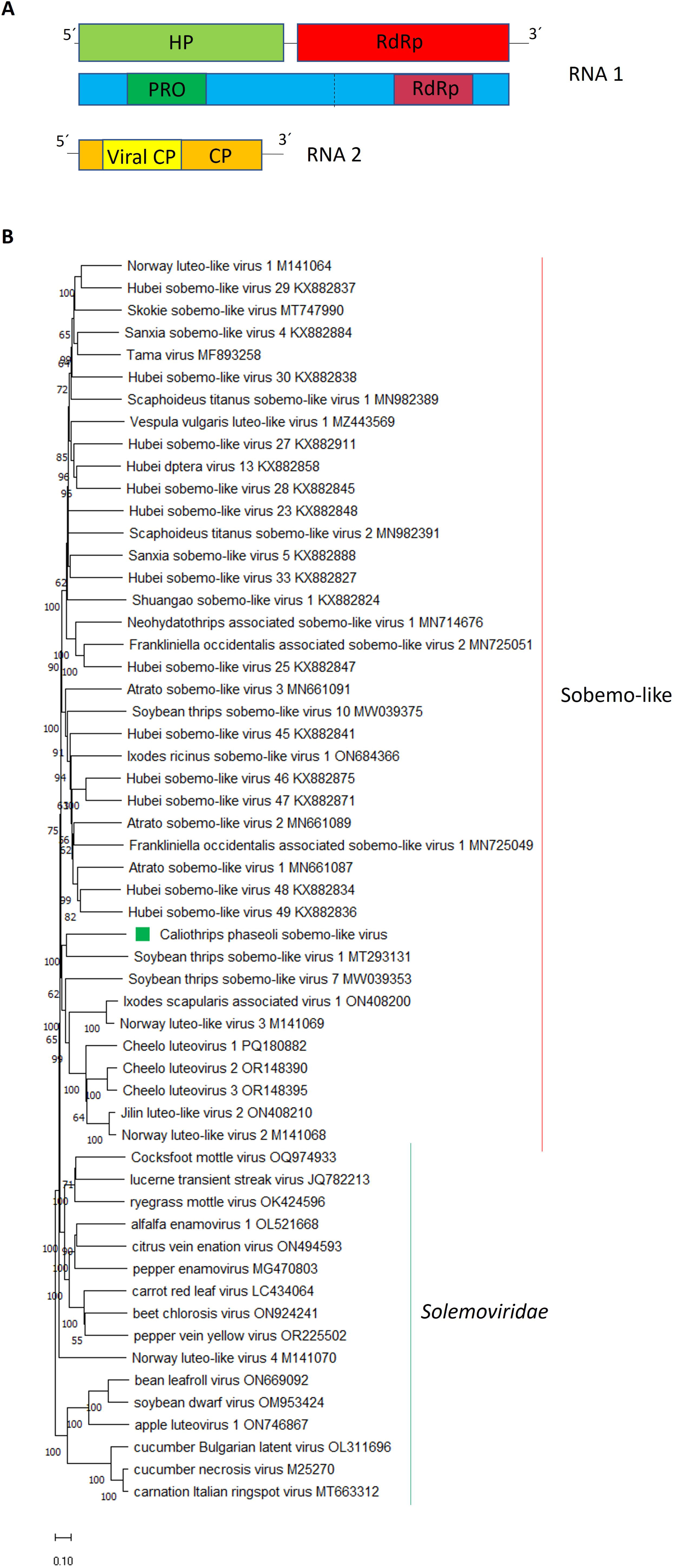
(A) Schematic representation of the genome organization of Caliothrips phaseoli sobemo-like virus (B) Maximum likelihood phylogenetic trees reconstructed using the RdRp sequence of Caliothrips phaseoli sobemo-like virus and of representative sobemo-like viruses. Bootstrap values above 50% are shown (1000 replicates). Caliothrips phaseoli sobemo-like virus is indicated with a green square. The scale bar shows the substitution per site.

Segment 1 of CPSlV contains two predicted ORFs, designated HP and RdRp. The HP ORF encodes a putative hypothetical protein (HP), whereas the second ORF encodes the RNA-dependent RNA polymerase (RdRp), which is expressed as a fusion polyprotein via a −1 ribosomal frameshifting event (Figure X). The HP protein has a length of 631 aa. Sequence comparisons using BlastP revealed that the CPSlV HP shares its highest similarity (27.86% identity) with the Soybean thrips sobemo-like virus 1 HP. The analysis of conserved domains identified a trypsin-like serine protease (PRO) domain located at aa positions 212-353 in the CPSlV encoded HP protein (Figure 2A). The RdRp protein has a size of 456 aa. BlastP analysis showed that the CPSlV RdRp is most closely related to Soybean thrips sobemo-like virus 1 RdRp, with 37.05% identity. A conserved viral RNA-dependent RNA polymerases (RdRp) domain was detected, which was located between the aa residues 153–364 (Figure 2A). The arrangement of genes within segment 1 of CPSlV resembles that one displayed by other arthropod-associated sobemo-like viruses (Otatti et al., 2020; Aragao et al., 2023; Dahan et al., 2024; Debat et al., 2025; Serra et al., 2026; Candia et al., 2026). This genomic organization looks like the 5′-proximal portion of genomes described for members of the *Solemoviridae* family, as the first ORF encodes a polyprotein that includes a serine protease domain, whereas the RdRp is synthesized from the downstream ORF through a −1 programmed ribosomal frameshifting (−1 PRF) mechanism (Debat et al., 2025; Candia et al., 2026). Nevertheless, in contrast to plant-infecting solemovirids, these viruses do not encode any ORF with a RNA silencing suppression function, consistent with their non-plant hosts (Debat et al., 2025; Candia et al., 2026).

Segment 2 of CPSlV contains a single ORF encoding a capsid protein (CP) (Figure 2A), with a length of 407 aa. Comparative BlastP analysis indicates that the CP of CPSlV shares its highest similarity (30.85% identity) with Frankliniella occidentalis associated sobemo-like virus 2 encoded CP. Domain analysis revealed the presence of a conserved viral coat protein (Viral CP) domain in the CP of CPSlV, which was located between the aa residues 65–243 in (Figure 2A).

Interestingly, any segment 2 was reported for the ten thrips-associated solemo-like viruses identified by Thekke-Veetil et al., (2020). Thus, it is tempting to speculate that authors did not search for that segment in their data.

These findings provide additional support for the hypothesis that the genomes of arthropod-associated sobemo-like viruses are bipartite; although earlier studies did not identified the second segments in many of these viruses, due to the uniqueness and highly divergence of this segment. Moreover, CPSlV CP, together with those reported from related arthropod-derived sobemo-like viruses, share recognizable amino acid similarity with capsid proteins characteristic of noda-like and permutotetra-like viruses. Such relationships are notable because, within those viral lineages, structural proteins are commonly expressed either from subgenomic RNAs or from independent genome segments. One possible explanation for these similarities is the occurrence of horizontal genetic exchange between distinct viruses. Such transfer events could result from erroneous encapsidation or segment exchange during mixed infections occurring within the same host species, thereby facilitating the movement of genomic material across unrelated viral taxa (Debat et al., 2025; Candia et al., 2026).

The phylogenetic analysis using the RdRp aa sequences located CPSlV within the lineage of unclassified solemovirid-related viruses detected in diverse arthropod hosts; where CPSlV clustered together with soybean thrips sobemo-like virus 1 (Figure 2B).

Taken together, both the phylogenetic evidence as well as the distinctive organization of their genomes, indicate that solemovirid-related viruses associated with non-plant hosts, especially arthropods, constitute a lineage that is clearly differentiated from currently recognized members of the order *Sobelivirales*. Their divergence may warrant the establishment of a separate viral family, within several genera should be assigned. Furthermore, the unusual genomic arrangements identified in these viruses suggest that they may employ replication and expression mechanisms that differ from those described for plant-infecting relatives. Elucidating these molecular strategies will be important not only for understanding their biology, but also for reconstructing the evolutionary events that contributed to the emergence and diversification of this viral clade.

#### 3.3.3. Molecular and phylogenetic characterization of the newly identified bean thrips Picorna-like virus

One contig shows similarity to picorna-like virus sequences. Nine recognized families were assigned to the viral order *Picornavirales*, where numerous representatives infect arthropod hosts (Zell et al., 2022). Members of this order have positive-sense single-stranded RNA genomes (Le Gall et al., 2008). Even though all picornavirids have a polyadenylated RNA, their display distinct genomic organizations; genomes may be either monopartite or bipartite and can encode between one and five open reading frames (Zell et al, 2022). Here, we identified a novel picorna-like virus, which we named as *Caliothrips phaseoli* picorna-like virus (CPPlV). The nearly complete genome of CPPlV comprises 9802 nt encoding a partial polyprotein (3’ end is missing) with a size of 2975 aa (Fig. 3A, Table 1).

**Figure 3.**
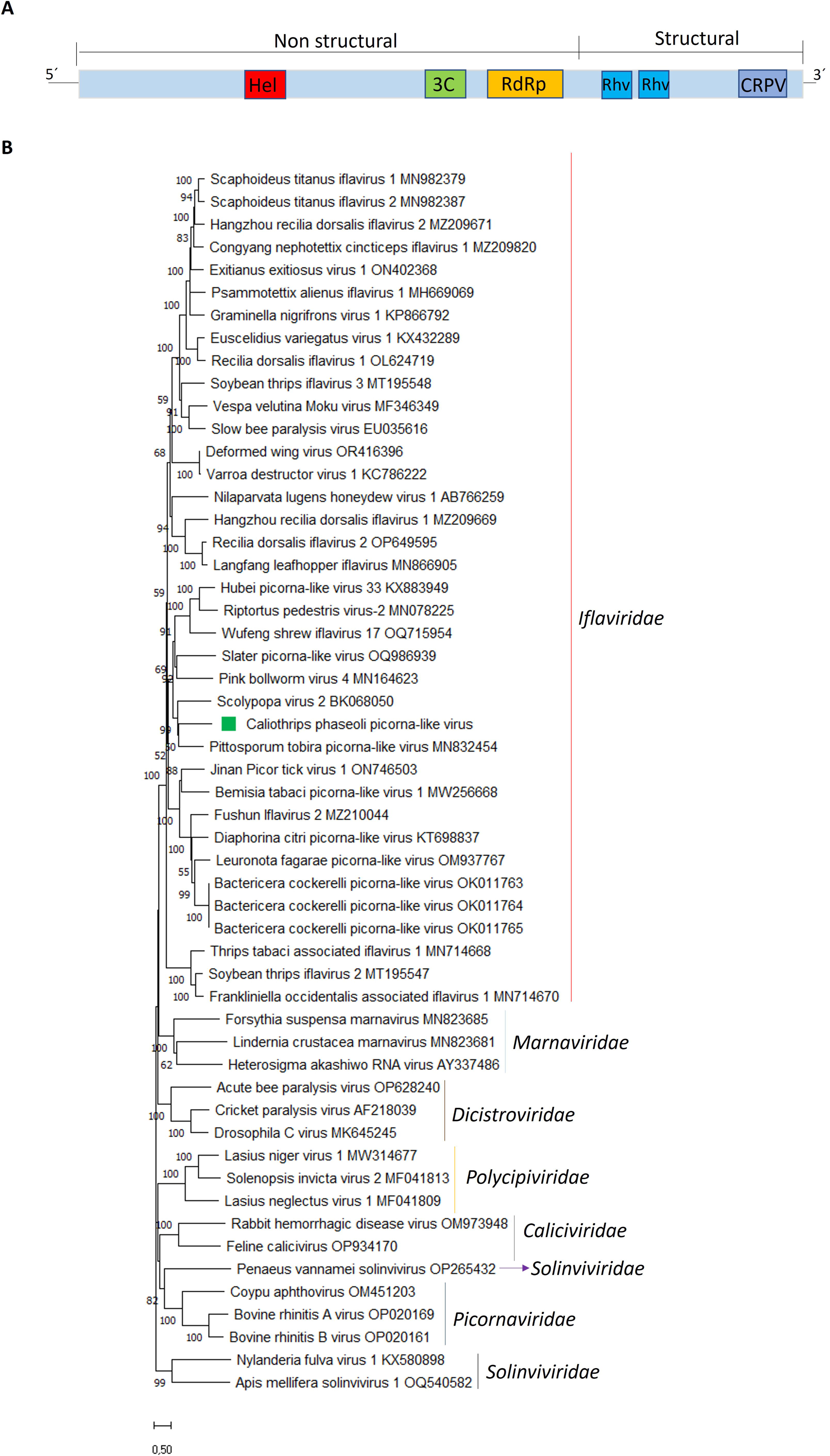
(A) Schematic representation of the genome organization of Caliothrips phaseoli picorna-like virus. (B) Maximum likelihood phylogenetic trees reconstructed using the polyprotein sequence of Caliothrips phaseoli picorna-like virus and of representative picorna-like viruses. Bootstrap values above 50% are shown (1000 replicates). Caliothrips phaseoli picorna-like virus is indicated with a green square. The scale bar shows the substitution per site.

Comparative BlastP analysis indicated that the CPPlV polyprotein is most closely related to that of the planthopper-associated Scolypopa virus 2 sharing 28.42% aa identity (Table 1). The genome layout of CPPlV is similar to that one described for other picorna-like viruses linked to distinct insect species which is named as type II organization (Zhang et al., 2023; Lahey et al., 2025), that exhibit a reversed genome organization in which those non-structural domains, which are involved in the replication, are positioned toward the N-terminus, whereas those structural domains are found in the C-terminal portion of the polyprotein (Figure 3A). However, those iflaviruses that were previously identified in thrips display the type I genome organization (Thekke-Veetil et al., 2020; Chiapello et al., 2021).

Within the N-terminal segment of the CPPlV polyprotein, the detected conserved domains were an RNA helicase (RNA_Helicase, aa 544-711), a Picornavirales 3C/3Clike protease (3C-Pro, aa 1238-1461), and a RNAdirected RNA polymerase (RdRp, aa 1499-1996) (Figure 3A). In contrast, the C-terminal region contains conserved domains associated with virion structure, specifically two rhinovirus-like capsid regions (Rhv-like, aa 2032-2283 and aa 2302-2577) as well as a cricket paralysis virus–like capsid domain (CRPV_capsid superfamily, aa 2752-2975) (Figure 3A). These conserved domains identified in the CPPlV polyprotein are characteristic of picornaviruses (Zell et al., 2022) supporting the placement of this newly identified virus within the order *Picornavirales*.

Phylogenetic analysis using the polyprotein sequence showed that CPPlV was clustered together with Scolypopa virus 2 and Pittoposporum tobira picorna-like virus which also display a type II genomic organization (Zhang et al, 2023; Lahey et al., 2025); and this clade is grouped with another cluster composed of picorna-like viruses that also have a type II genomic organization (Fig 3B) (Zhang et al., 2023). This clustering indicates that these viruses that share a same genomic organization likely originated from a common ancestor characterized by an inverted genome organization relative to that of members of the family *Iflaviridae*. Although CPPlV and related viruses genomes shows a reversed layout and share a low levels of identity between them, they are clearly grouped with members belonging to the family *Iflaviridae* (Figure 3B). However, CPPlV and related viruses that share a common evolutionary history should be assigned to a novel genus within this family. The further discovery of novel members exhibiting a type II genomic organization related to CPPLV and its clade will provide further evidence in favor of formal recognition of a novel genus within the Iflaviridae family by the ICTV.

Several picornaviruses have been reported to exert detrimental effects on their insect hosts, including developmental deformities, behavioral alterations, and, in certain instances, death (Mattia et al., 2026; Li et al., 2026). Consequently, further research is needed to determine whether the recently discovered picorna-like virus detected in bean thrips is capable of inducing pathogenic responses in this insect species. Evaluating these host-virus interactions may provide valuable insight into the feasibility of exploiting this virus as a biological control strategy to reduce thrips infestations in peanut and soybean plantations.

#### 3.3.4. Molecular and phylogenetic characterization of the newly identified bean thrips virga-like virus

One contig shows similarity to virga-like virus sequences. Virgavirids have positive-sense single-stranded RNA genomes (Kondo et al., 2019). Here, we identified a novel virga-like virus, which we named as *Caliothrips phaseoli* virga-like virus (CPVlV).

The coding-complete genome of CPVlV comprises 11866 nt and has three ORFs, where ORF1 encodes a polyprotein with a size of 2623 aa, ORF2 encodes the RdRp with a size of 581aa, while ORF3 encodes a putative CP with a size of 140aa (Fig. 4A, Table 1).

**Figure 4.**
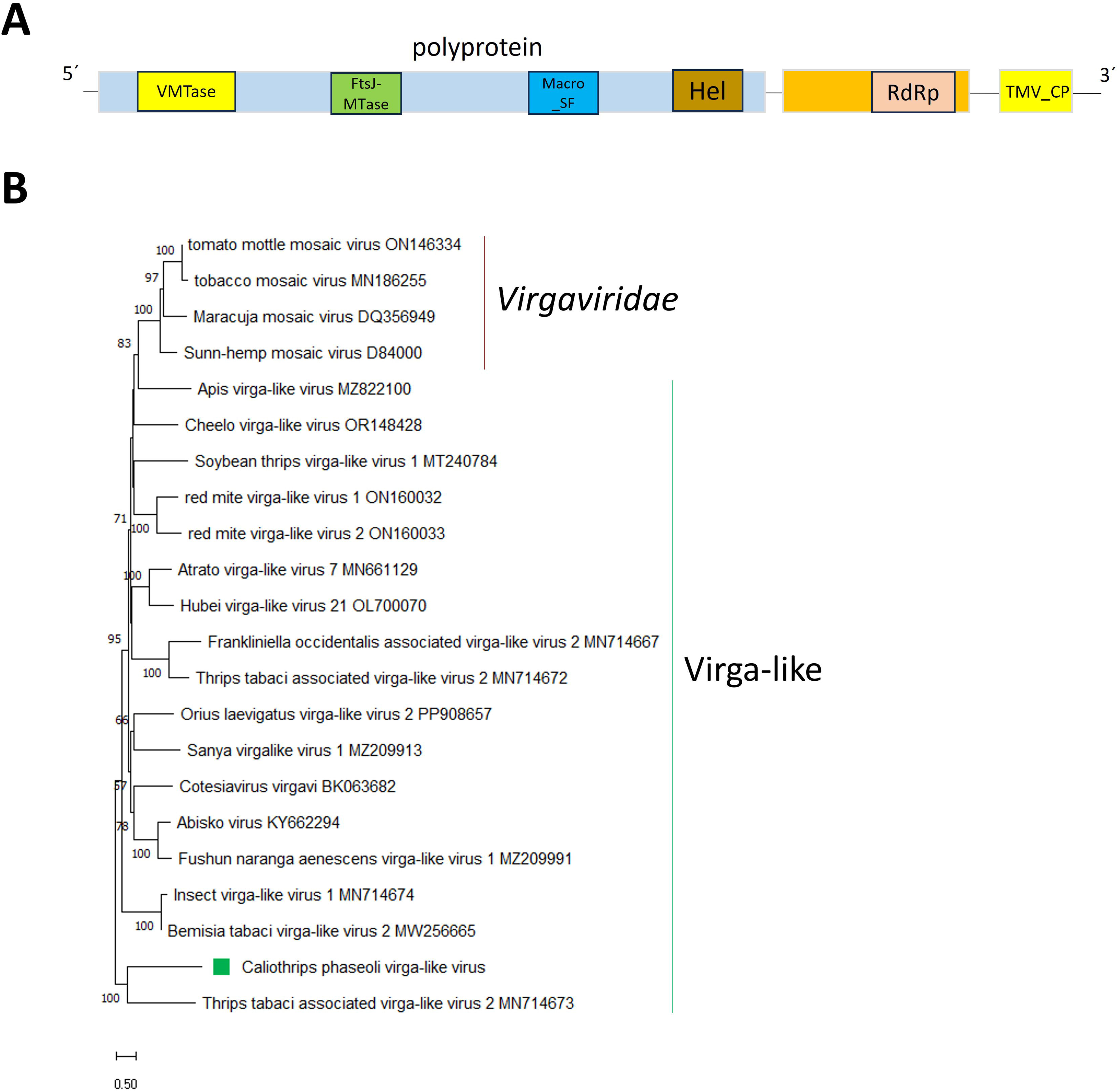
(A) Schematic representation of the genome organization of Caliothrips phaseoli virga-like virus. (B) Maximum likelihood phylogenetic trees reconstructed using the RdRp sequence of Caliothrips phaseoli virga-like virus and of representative virga-like viruses. Bootstrap values above 50% are shown (1000 replicates). Caliothrips phaseoli virga-like virus is indicated with a green square. The scale bar shows the substitution per site.

Comparative BlastP analysis indicated that the CPVlV polyprotein is most closely related to that of the thrips-associated Thrips tabaci-associated virga-like virus 2 (TTaVlV2) sharing 28.52% aa identity, while the CPVlV RdRp is most closely related to that of spider virga-like virus 59 sharing 38.30% aa identity; and the CPVlV CP is most closely related to that of Drosophila crammond virga-like virus sharing 29.57% aa identity (Table 1). The genome layout of CPVlV is similar to that one described for the thrips associated Frankliniella occidentalis associated virga-like virus 2 (Chiapello et al., 2021).

Within the N-terminal segment of the CPVlV polyprotein, the detected conserved domains were a Viral methyltransferasean (VMTase, aa 173-425) and a FtsJ-like methyltransferase (FtsJ, aa 938-1128); while the conserved domains identified in the C-terminal region of the encoded polyprotein were a macrodomain, Poa1p-like family (Macro_Poa1p-like, aa 1798-1928) and a Viral (Superfamily 1) RNA helicase (Viral_Helicase1, aa 2323-2574). The conserved domain ps-ssRNAv_RdRp-like Superfamily (RdRp, aa 248-511) was identified in the CPVlV RdRp, while the CPVlV CP has a TMV-like viral coat proteins (TMV_coat, aa 3-139) (Figure 4A). These conserved domains identified in the CPPlV are characteristic of several virga-like viruses associated to insects (Chiapello et al., 2021) supporting the placement of this newly identified virus within this group of viruses.

Phylogenetic analysis using the polyprotein sequence showed that CPVlV was clustered together with the thrips associated TTaVlV2; and this clade is grouped with another cluster composed of virga-like viruses (Fig. 4B). However, identity values as well as evolutionary relationships shows how distinct are CPVlV and TTaVlV2, and these viruses from other insect-associated viurga-like viruses, illustrating that a huge amount of viral “dark matter” is present in virga-like viruses associated to insects. Virga-like viruses fill the evolutionary gap between insect-specific negeviruses and plant specific virgaviruses (Kondo et al., 2019) Thus, virga-like viruses associated to insects should likely be accommodated in a new family with several genera to be assigned within the order *Martellivirales*.

### 3.4. (-)ssRNA viruses

### 3.4.1. Molecular and phylogenetic characterization of the newly identified bean thrips thogotovirus

Five contigs show similarity to thogotovirus sequences. *Thogotovirus* is a genus belonging to the family *Orthomyxoviridae*. Members of this genus are enveloped and have a single-stranded, negative-sense RNA multi segmented genome with five to seven segments, and are generally tick-associated (Lambert et al., 2015), although several of them have also been identified in insects (Da Silva et al., 2023; Petrone et al., 2023), including thrips (Thekee-Veetil et al., 2020).

Here, a novel thogotovirus was identified and named as Caliothrips phaseoli thogotovirus (CPTV) The CPTV genome sequence comprised five segments of single-stranded, negative-sense RNAs of 2431 nt, 2207 nt, 2021 nt, 1698 nt, and 1553 nt (Table 1). Each segment has a single ORF. Segment 1 encodes the PB2 polymerase, that have a 758 aa size. PB2 protein shows the highest BlastP similarity with soybean thrips thogotovirus 1 (STTV1)-PB2, with 46.67% identity and a Flu_PB2 conserved domain has been identified in its sequence at aa positions 7-440. Segment 2 encodes the PB1 polymerase, that have a 713 aa size. PB1 protein shows the highest BlastP similarity with STTV1-PB1, with 61.34% identity and a Flu_PB1 conserved domain has been identified in its sequence at aa positions 6-675. Segment 3 encodes the PA polymerase, that have a 649 aa size. PA protein shows the highest BlastP similarity with STTV1-PA, with 51.4% identity and a Flu_PA conserved domain has been identified in its sequence at aa positions 30-644. Segment 4 encodes a glycoprotein (GP), that have a 508 aa size. The GP shows the highest BlastP similarity with STTV1-GP, with 40.89% identity and an Env_GP conserved domain has been identified in its sequence at aa positions 32-504. Segment 5 encodes a nucleoprotein (NP), that have a 458 aa size. The NP shows the highest BlastP similarity with STTV1-NP, with 40.34% identity and a Flu_NP-like conserved domain has been identified in its sequence at aa positions 184-390 (Table 1, Fig. 5A).

**Figure 5.**
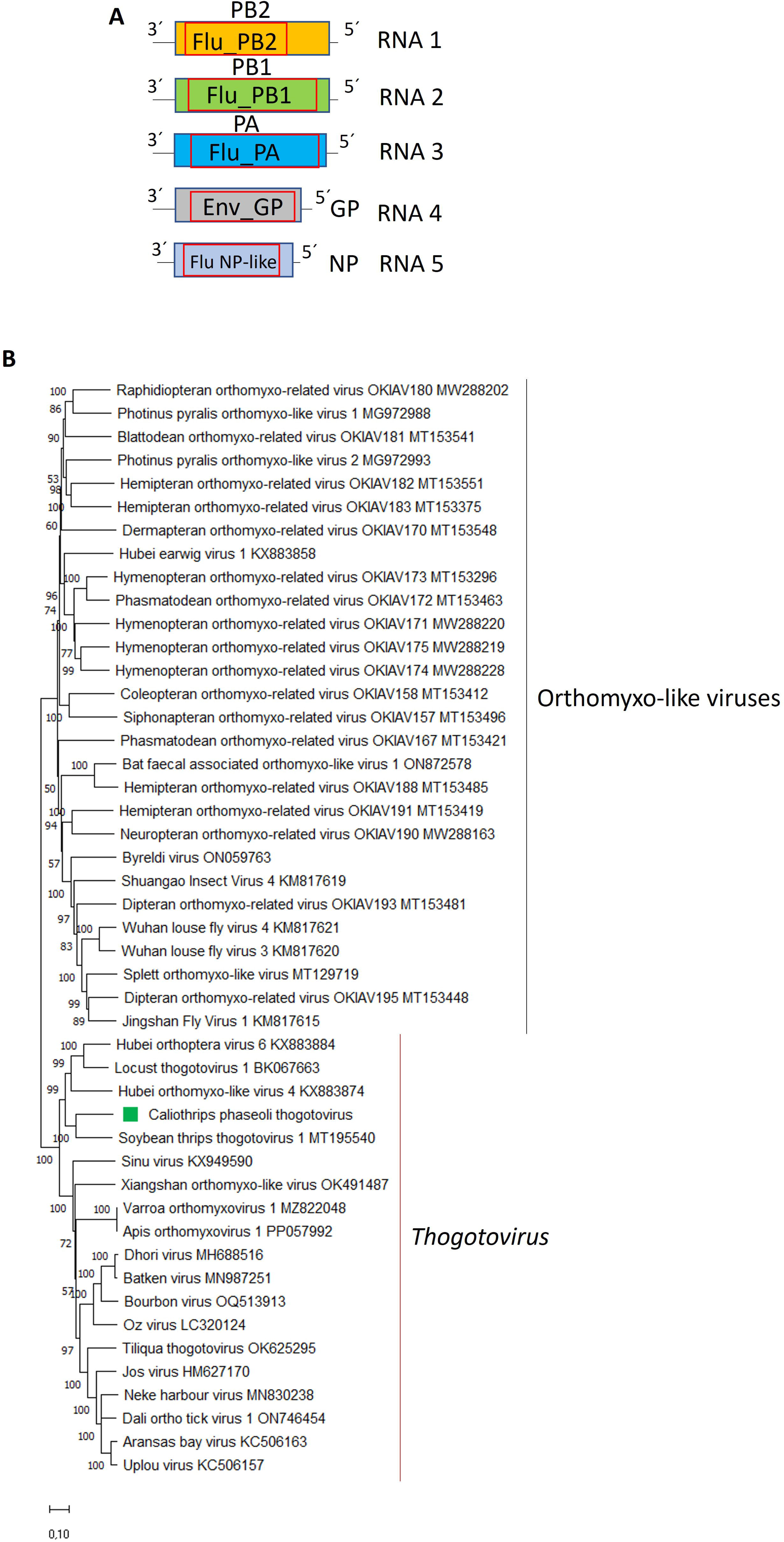
(A) Schematic representation of the genome organization of Caliothrips phaseoli thogotovirus. (B) Maximum likelihood phylogenetic trees reconstructed using the PB1 sequence of Caliothrips phaseoli thogotovirus and of representative thogotoviruses and orthomyxo-like viruses. Bootstrap values above 50% are shown (1000 replicates). Caliothrips phaseoli thogotovirus is indicated with a green square. The scale bar shows the substitution per site.

The phylogenetic analysis based on the PB1 protein, placed CPTV in a cluster with STTV1 (Fig. 5B), which also has 5 segments in its genome (Thekke-Veetil et al., 2020) and this cluster is related to a clade formed by Hubei orthoptera virus 6, that also has a genome composed of 5 segments (Shi et al., 2016), Hubei orthomyxo-like virus 4 and Locus thogotovirus 1 (Fig. 5B), which only one segment was identified to date (Shi et al., 2016; Petrone et al., 2023). It is tempting to speculate that this five viruses share a common ancestor which adapted to infect insects and the segment encoding the matrix proteins was lost during its evolution. Moreover, these viruses, which appear to be prevalent on insects (Fallon et al, 2018), could be accommodated in a novel genus within the family *Orthomyxoviridae* rather than as members of the genus *Thogotovirus*.

#### 3.4.2. Molecular and phylogenetic characterization of the newly identified bean thrips phenuivirus

Six contigs showed similarity to sequences from members of the family *Phenuiviridae*, order *Hareavirales*, which includes viruses that infect a wide range of hosts, including human, mammals, invertebrates, and plants (Sasaya et al., 2023). This family is composed of viruses with two to eight segments of negative-sense or ambisense RNA (Sasaya et al., 2023). The three identified phenuiviruses were named as Caliothrips phaseoli phenuivirus 1-3 (CPPV1-3) (Table 1).

For each one of the three viruses, two segments were assembled (Table 1, Fig. 6A-C). For CPPV1, segment 1 has a sequence length of 7160 nt, while segment 2 is composed of 3365 nt. For CPPV2, segment 1 has a sequence length of 6464 nt, while segment 2 is composed of 2217 nt. While for CPPV3, segment 1 has a sequence length of 7258 nt, while segment 2 is composed of 1465 nt (Table 1).

**Figure 6.**
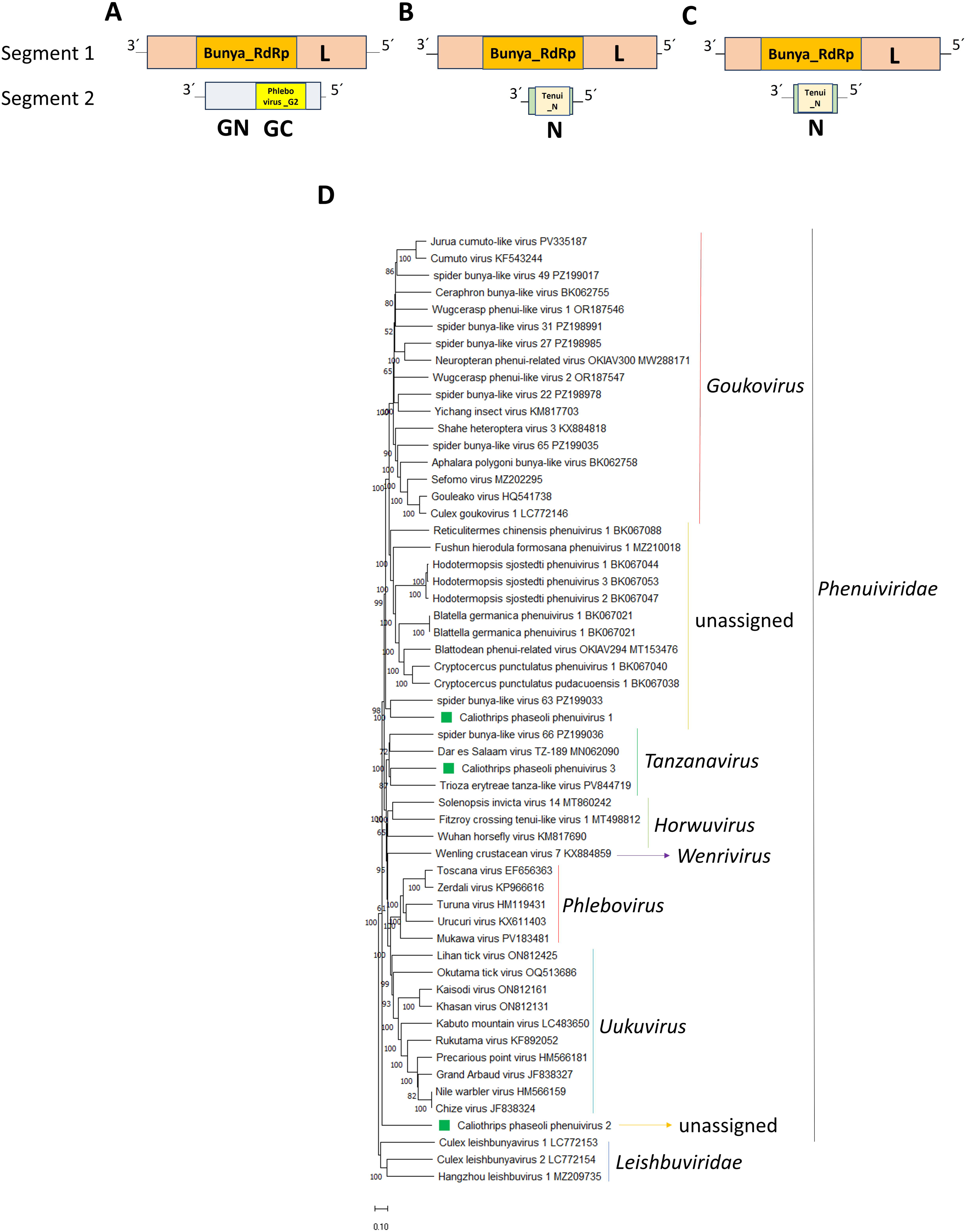
(A) Schematic representation of the genome organization of Caliothrips phaseoli phenuivirus 1-3. (B) Maximum likelihood phylogenetic trees reconstructed using the RdRp sequence of Caliothrips phaseoli phenuivirus 1-3 and of representative phenuiviruses. Bootstrap values above 50% are shown (1000 replicates). Caliothrips phaseoli phenuivirus 1-3 are indicated with a green square. The scale bar shows the substitution per site.

Segment 1 of CPPV1-3 contains a single ORF encoding an RdRp with a size of 2298 aa, 2111aa and 2315aa, respectively (Fig. 6A). BLASTp searches of the nr protein database using the CPPV1-3 RdRp showed that the best hits were the RdRp encoded by spider bunya-like virus 63, spider bunya-like virus 27 and Trioza eritreae tanza-like virus, with identities values of 34.30, 26.32 and 32.38%, respectively (Table 1). The conserved superfamnily domain Bunya_RdRp was identified in CPPV1-3 RdRp’s (Fig. 6A-C), which is the commonly motif found in the phenuivirids (Sasaya et al., 2023).

The second segment assembled for CPPV1 contains a single ORF encoding a glycoprotein precursor (GP) of 1027 aa (Table 1, Fig. 6A). BLASTp searches of the nr protein database using the CPPV1 GP showed that the best hit was the GP encoded by Ceraphron bunya-like virus, with an identity value of 21.58% (Table 1). The conserved Phlebovirus glycoprotein G2 fusion domain (Phlebovirus_G2) was identified in the C-terminal region of the CPPV1 GP (Fig. 6A). This conserved domain is commonly found in phenuivirids and is a critical structural region which acts as a class II viral fusion protein (Spiegel et al., 2016).

The second segment assembled for CPPV2-3 contains a single ORF encoding a nucleoprotein (NP) of 615 aa and 317 aa, respectively (Table 1, Fig. 6B-C). BLASTp searches of the nr protein database using the CPPV2-3 NP showed that the best hits were the NP encoded by Okutama tick virus and Trioza eritreae tanza-like virus, with identities values of 27.16 and 27.78%, respectively (Table X). The Tenui_N conserved domain was identified in the CPPV2-3 NPs (Fig. 6B-C). This domain is commonly found in phenuivirids (Sasaya et al., 2023).

Interestingly any segment encoding the NP was found for CPPV1 and any segment encoding the GP was found neither in CPPV2 nor CPPV3 (Table 1, Fig. 6A-C). Thus, it is highly likely that the genome descriptions of CPPV1-3 are incomplete because of our inability to recover a third segment for each one of these viruses due to the high divergence shared with other members.

The phylogenetic tree based on RdRp aa sequence alignment placed CPPV1, CPPV2 and CPPV3 with other phenuivirids in different clusters (Fig. 6D). CPPV1 clustered with spider bunya-like virus 63, forming a clade related to other unassigned phenuivirids (Fig. 6D). CPPV2 is placed in a monophyletic cluster distantly related to the cluster grouping phenuivirids (Table 6D); while CPPV3 is grouped together with Trioza eritreae tanza-like virus in a clade grouping members belonging to the genus *Tanzanavirus*, family *Phenuiviridae* (Fig. 6D). CPPV1, CPPV2 and CPPV3 represent novel phenuivirus members, where CPPV1 and CPPV2 should be assigned to two newly proposed genera to be created within the family *Phenuiviridae;* while CPPV3 shoulkd be assigned as a novel member within the genus *Tanzanavirus*.

### 3.5. dsRNA viruses

#### 3.5.1. Molecular and phylogenetic characterization of the newly identified bean thrips partitiviruses

Nine contigs show similarity to partitivirus sequences. The nine identified segments correspond to four putative viruses, which were named Caliothrips phaseoli partitivirus 1-4 (CPPtV1-4) (Table 1). The members formally assigned to the family *Partitiviridae* by the ICTV have monocistronic, bi-segmented dsRNA genomes where the firs segment encodes the RdRp, while the second segment encodes the CP (Vainio et al., 2018); however, some unassigned partitiviruses have multisegmented genomes composed of three, four or even five segments (Thekee-Veetil et al., 2020; Ma et al., 2024; Liu et al., 2025; Zhou et al., 2025). Assigned partitiviruses have plants or fungi as their hosts (Vainio et al., 2018); however, partiti-like viruses have been identified in insect (Shi et al., 2016; Cross et al., 2020; Wu et al., 2020; Xu et al., 2020), including thrips (Thekee-Veetil et al., 2020; Chiapello et al., 2021).

The full-length coding region of CPPtV1-3 RNA1 has a length of 2298 nt, 1858nt and 1532 nt, respectively (Table 1), and encodes a putative RdRp (Fig. 7A-B) of 715 aa, 575aa and 464 aa, respectively. BLASTp searches showed that the CPPtV1-3 RdRp best hit was the RdRp protein encoded by Plasmopara viticola lesion-associated Partitivirus 7 (PVaPV7), with a 50.14% identity, the Aspergillus flavus partitivirus 2 RdRp with a 53.45% identity and the RdRp protein encoded by Scaphoideus titanus-associated partiti-like virus 1 (STaPlV1) with a 52.05% identity, respectively (Table 1). For CPPtV4 the RNA segment encoding the putative RdRp was partially assembled (Table 1). BLASTp searches of the nr protein database showed that the best hit was the RdRp protein encoded by STaPlV1 (Table 1). The conserved domain RdRp_1 was identified in the CPPtV1-4 RdRps (Fig. 7A-B).

**Figure 7.**
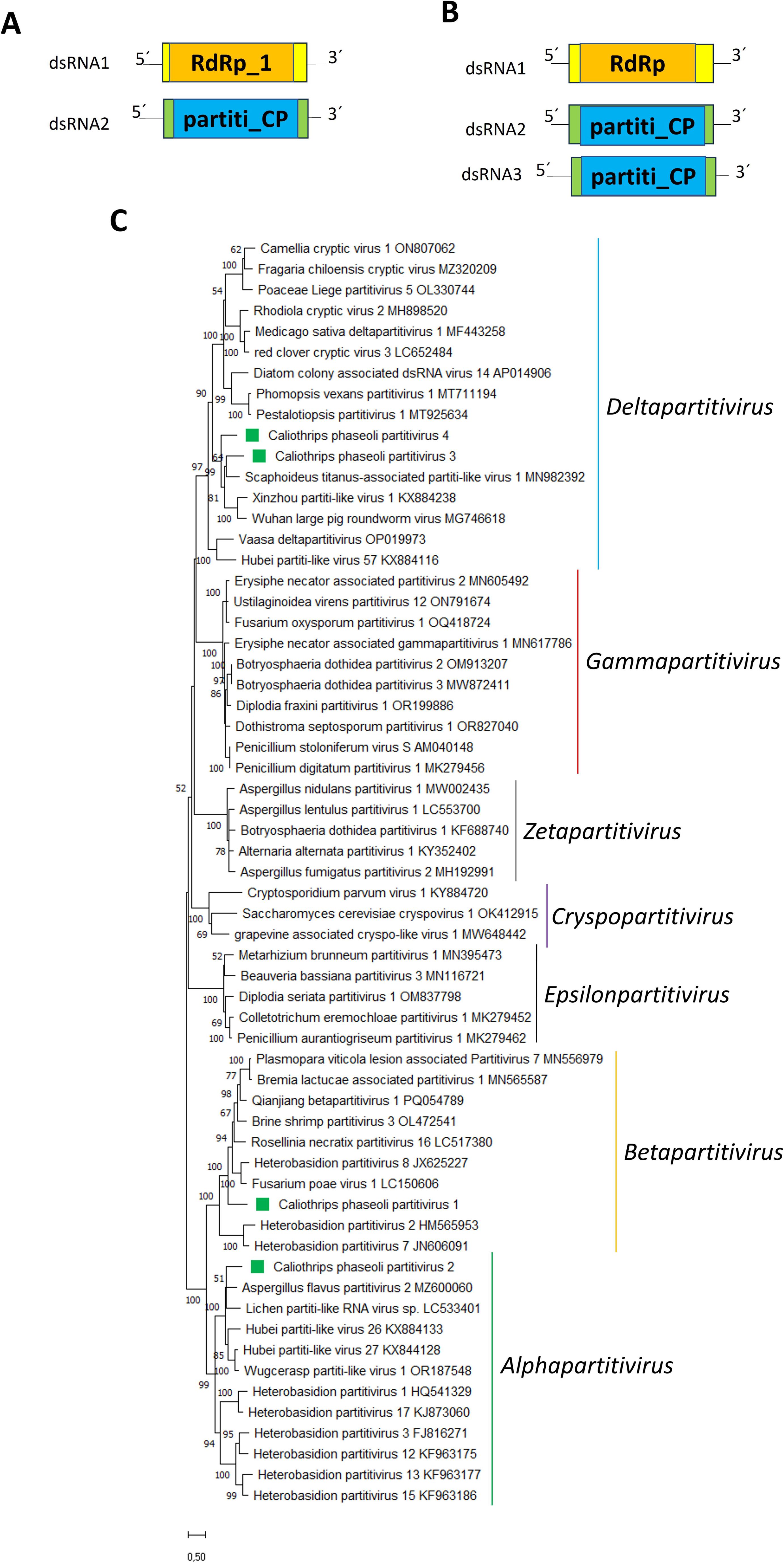
(A) Schematic representation of the genome organization of Caliothrips phaseoli partitivirus 1-4. (B) Maximum likelihood phylogenetic trees reconstructed using the RdRp sequence of Caliothrips phaseoli partitivirus 1-4 and of representative partitiviruses. Bootstrap values above 50% are shown (1000 replicates). Caliothrips phaseoli partitivirus 1-4 are indicated with a green square. The scale bar shows the substitution per site.

The full-length coding region of CPPtV1-4 RNA2 has a length of 2282 nt, 1759nt, 1329nt and 1367nt, respectively (Table 1), and encodes a putative CP of 657aa, 507aa, 325aa, and 301aa, respectively (Table 1). BLASTp searches showed that the CPPtV1-4 CP best hit was the CP encoded by PVaPV7, Lichen partiti-like virus (LiPlV), STaPlV1 and STaPlV1 with a 36.64%, 29.31%, 35.46% and 40.69% identity, respectively (Table 1). The conserved domain partiti_CP was identified in the CPPtV1-4 CPs (Fig. 7A-B).

Interestingly a third segment was assembled for CPPtV2 (Fig. 7B). The full-length coding region of this segment has 1756nt in length and encodes a putative CP of 467aa, which best hit is the LiPlV encoded CP with a 24.58% identity (Table 1), and the conserved domain partiti_CP was identified in its sequence (Table 1). The fungi-associated partiti-like virus Cladosporium cladosporioides partitivirus 2 also has three segments, with two of them encoding putative CPs (Jaccard et al., 2023)

The phylogenetic tree based on RdRp aa sequence alignment placed CPPtV1 with other betapartitiviruses (Fig. 7C), CPPtV2 was placed in another clade together with other alphapartitiviruses, including Aspergillus flavus partitivirus 2 and Lichen partiti-like virus (Fig. 7C); whose genomes also have more than two segments (Urayama et al., 2020; Degola et al, 2021) such as CPPtV3. On the other hand, CPPtV3 and 4 clusters with other deltapartitiviruses (Fig. 7C).

Members of the genera *Alphapartitivirus*, *Betapartitivirus* and *Deltapartitivirus* infect fungi or plants. (Vainio et al., 2018). Thus, it is tempting to speculate that CPPtV1-4 are mycoviruses that could be related to the presence of fungi being part of the thrips-associated microbiome; however, since some partiti-like viruses can infect insects (Cross et al., 2020; Xu et al., 2020), there is a possibility that CPPtV1-4, or some of them, are indeed true thrips viruses. Recently, it has been shown that alpha- and betapartitiviruses were able to replicate in insects (Telengech et al., 2024), supporting the existence of true insect partitiviruses.

#### 3.5.2. Molecular and phylogenetic characterization of the newly identified bean thrips toti-like viruses

Four contigs show similarity to toti-like virus sequences. The four identified segments correspond to four putative viruses, which were named Caliothrips phaseoli toti-like virus 1-4(CPTlV1-4) (Table 1). Totiviruses have dsRNA genomes where most of its members have two ORFs (Hillman and Cohen, 2021), and several of them were identified in thrips (Thekke-Veetil et al., 2020; Chiapello et al., 2021; Zhang et al., 2025). Recently the taxonomy among totiviruses was reorganized to classify toti-like viruses into distinct newly created families (Sato et al., 2023), one of this newly created families is Spiciviridae, whose members have been identified in a remarkably broad range of hosts (do Socorro Foro Ramos et al., 2026). Spiciviruses have a non-segmented dsRNA genome with two open reading frames (ORFs) encoding a proline-alanine-rich protein (PArP) and an RNA-dependent RNA polymerase (RdRp) and these viruses may lack a capsid, an atypical trait among dsRNA viruses (do Socorro Foro Ramos et al., 2026). Moreover, it has been proposed that the unique combination of broad host range and distinct genomic architecture positions Spiciviridae as a key group for studying RNA virus evolution in arthropods (do Socorro Foro Ramos et al., 2026).

The full-length RdRp coding region was assembled for CPTlV1 and 2; where CPTlV1-2 assembled fragments have a length of 4525 nt, and 4864 nt, respectively (Table 1), and encodes a putative RdRp (Fig. 8A) of 1321 aa, and 1378 aa, respectively. BLASTp searches showed that the CPPtV1-2 RdRp best hit was the RdRp protein encoded by Spider totivirus 76 (PVaPV7), with a 34.46% identity, and the RdRp protein encoded by Spider totivirus 1 with a 32.02% identity, respectively (Table 1). On the other hand, two ORFs were identified in the partial genomes of 6512nt and 5875nt assembled for CPTlV3-4, respectively (Table 1; Fig. 8B). CPTlV3-4 ORF1 encode a putative PArP, partially assembled, which best hits were PArP protein encoded by Spider totivirus 33 and Spider totivirus 99, with identities below 33% (Table 1). Any conserved domain was identified in the CPTlV3-4 PArP proteins (Fig. 8B). CPTlV3-4 ORF2 encodes the RdRP, which was fully assembled for CPTlV3 with a length of 1278 aa and partially assembled for CPTlV4 (Table 1) he RNA segment encoding the putative RdRp was partially assembled (Table 1). BLASTp searches of the nr protein database showed that the best hit for CPTlV3 were the RdRp protein encoded by Spider totivirus 33 and, respectively, with identities below 44% (Table 1). The conserved domain RdRp was identified in the CPTlV1-4 RdRps (Fig. 8A-B).

**Figure 8.**
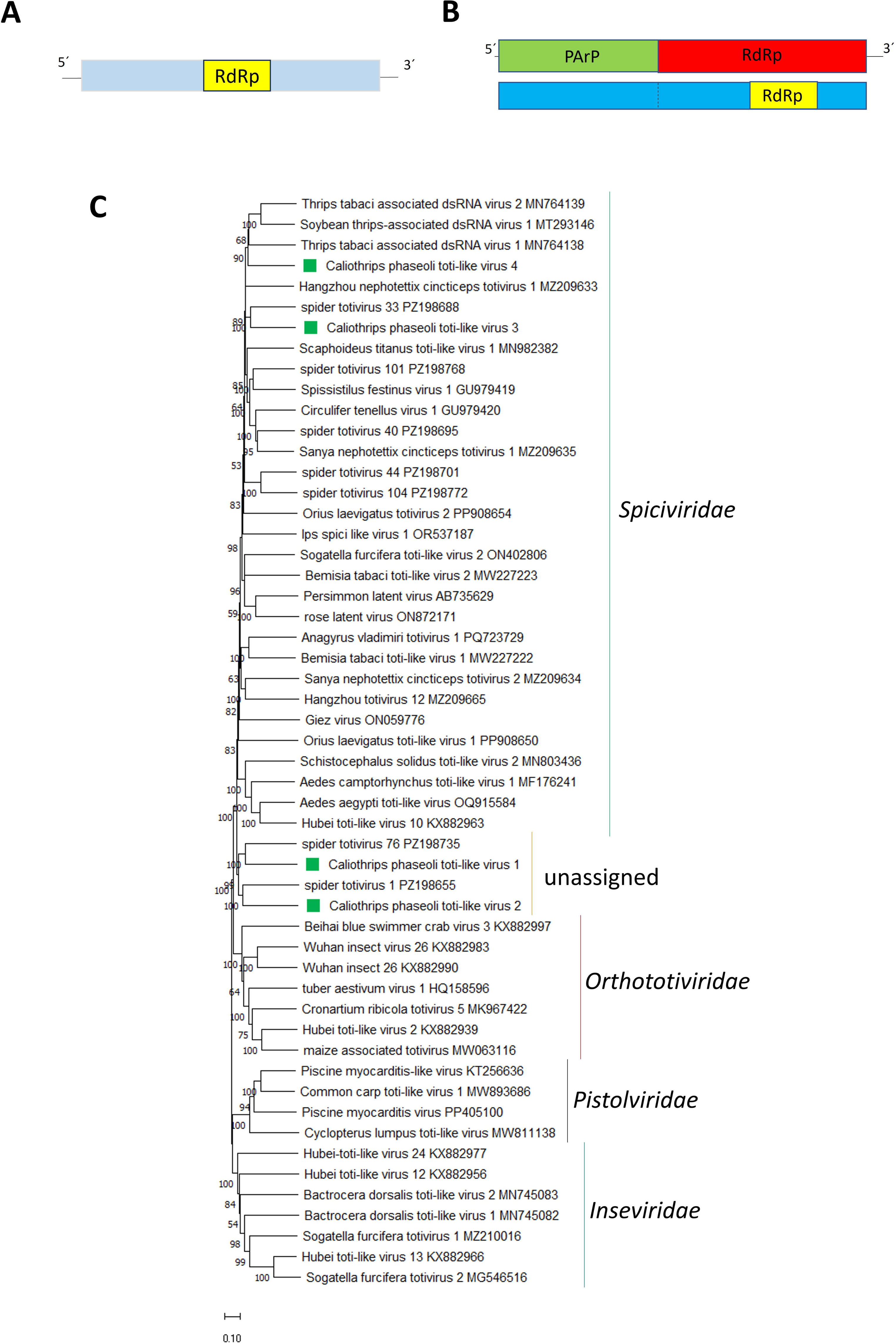
(A) Schematic representation of the genome organization of Caliothrips phaseoli toti-like virus 1-4 (B) Maximum likelihood phylogenetic trees reconstructed using the RdRp sequence of Caliothrips phaseoli toti-like virus 1-4 and of representative toti-like viruses. Bootstrap values above 50% are shown (1000 replicates). Caliothrips phaseoli toti-like virus 1-4 are indicated with a green square. The scale bar shows the substitution per site.

The phylogenetic tree based on RdRp aa sequence alignment placed CPTlV3-4 with other spicivirids (Fig. 8C), while CPTlV1-2 were placed in a clade together with Spider totivirus 76 and Spider totivirus 1, outside of the spicivirids clade (Fig. 8C); thus CPTlV1-2 could likely be assigned to a distinct family rather than the Spiciviridae, the inability of find a second ORF for CPTlV1-2, as well as for Spider totivirus 76 and 1, supports this. However, the high divergence between these viruses and related ones, could be the reason of our inability to not identify a second ORF for CPTlV1-2. Moreover, the high divergence found illustrate that a huge amount of viral “dark matter” is still present in toti-like viruses associated to insects.

### 3.6. The bacterial microbiome carries a genuine insect-endosymbiont signal

The bacterial fraction is small in reads (183 rPM total) but structured like an insect microbiome (Figure S1): Pseudomonadota dominate both reads and richness (317 species, 93 rPM), followed by Bacteroidota (89 species, 33 rPM), Bacillota (122 species, 25 rPM) and Actinomycetota (68 species, 21 rPM). Within this background, recognized insect endosymbionts stand out: *Rickettsia bellii* (98.6% identity, 51 reads), *Rickettsia* sp. MEAM1 (96.9%) and *Rickettsia tamurae* (98.0%); three *Cardinium* endosymbionts (∼92%, 23–27 reads each) together with the *Cardinium*-related *Amoebophilus*; *Arsenophonus* endosymbiont of *Bemisia tabaci* (100%, 4 reads); and *Wolbachia pipientis* (100%, low read support) (Figure S2).

*Rickettsia*, *Cardinium*, *Arsenophonus* and *Wolbachia* have both the read support and the host-association precedent to be credible. *Cardinium* and *Wolbachia* co-infections are documented in haplodiploid thrips, where they independently induce cytoplasmic incompatibility (Nguyen et al., 2017) and influence host fitness and sex allocation (Katlav et al., 2022); *Rickettsia* is a widespread facultative endosymbiont of sap-feeding pests, as shown for whiteflies sharing the R*ickettsia*/*Arsenophonus*/*Cardinium* guild that we also detect here (Rossitto De Marchi and Smith, 2020). Thus, the presence of *Rickettsia bellii*, *Cardinium*, *Arsenophonus* and *Wolbachia* in our data is therefore biologically expected.

## Concluding remarks

The present study identified 18 previously undescribed viruses associated with the bean thrips through transcriptome-based sequencing, considerably expanding the known virome of this economically important insect. Beyond increasing the catalog of viruses infecting *Caliothrips phaseoli*, these discoveries provide valuable information for understanding the evolutionary history and diversity of viruses associated with thrips hosts. Collectively, the results demonstrate the effectiveness of high-throughput metatranscriptomic approaches for uncovering hidden viral diversity that would likely remain undetected using conventional methods.

An important next step will be to determine the biological consequences of these viral infections for their insect host. Functional studies examining virus–host interactions will help clarify whether these viruses influence key aspects of thrips biology, including development, reproduction, survival, behavior, immune responses, or adaptation to environmental stress. Such knowledge will improve our understanding of the ecological roles of insect-associated viruses and their contribution to host fitness and population dynamics.

From an applied perspective, these findings may also support the development of sustainable approaches for managing peanut thrips populations. Elucidating the mechanisms through which these viruses interact with their host could reveal opportunities to exploit naturally occurring viral processes as components of integrated pest management programs. Furthermore, establishing a comprehensive characterization of the *C. phaseoli* virome represents a critical step toward evaluating the feasibility of virus-based biological control agents. The viral resources described here therefore provide a valuable framework for future research aimed at designing environmentally friendly virocontrol strategies for this agriculturally important pest.

## Author Contributions

**NB**: conceptualization, investigation, formal analysis, funding acquisition, writing-original draft and editing. **SdB**: investigation, funding acquisition, writing-review and editing. **CD**: investigation, formal analysis, editing. **HD**: investigation, formal analysis, data curation, writing-review and editing. All authors have read and agreed to the published version of the manuscript.

## Institutional Review Board Statement

This article does not contain any studies with human participants or animals performed by any of the authors.

## Informed Consent Statement

Not applicable.

## Data Availability Statement

The sequences were deposited in GenBank with accession numbers

