## Supplementary material for "Shedding light on the virome of *Caliothrips phaseoli*, an emergent pest in peanut": Supp Table S1

Table S1. Peanuts fields evaluated

| Field ID | Growing season | Location (latitude and longitude) |
| --- | --- | --- |
| 1 | 2023-2024 | -31.745848; -63.781707 |
| 2 | 2023-2024 | -32.355183; -63.833729 |
| 3 | 2023-2024 | -34.814233; -64.152723 |
| 4 | 2023-2024 | -34.823608; -64.240954 |
