## Supplementary material for "Shedding light on the virome of *Caliothrips phaseoli*, an emergent pest in peanut": Supp Figure 1

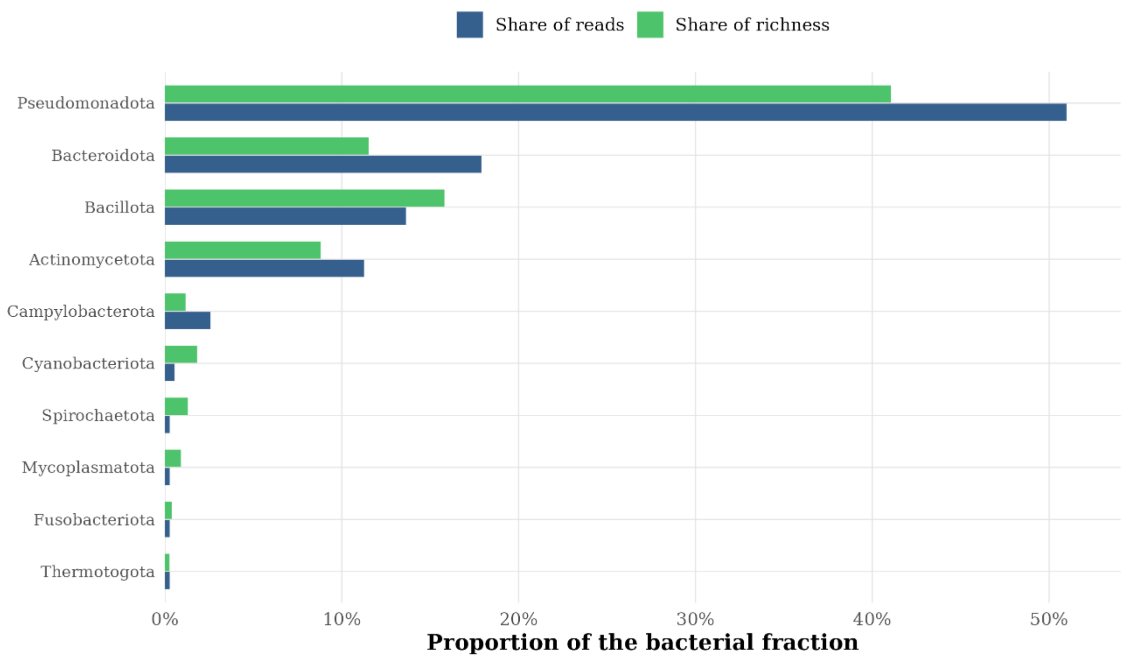


**Figure S1**. Bacterial microbiome at phylum level. For the ten most abundant bacterial phyla, the share of bacterial reads and the share of bacterial richness are shown side by side.
