## Supplementary material for "Shedding light on the virome of *Caliothrips phaseoli*, an emergent pest in peanut": Supp Figure 2

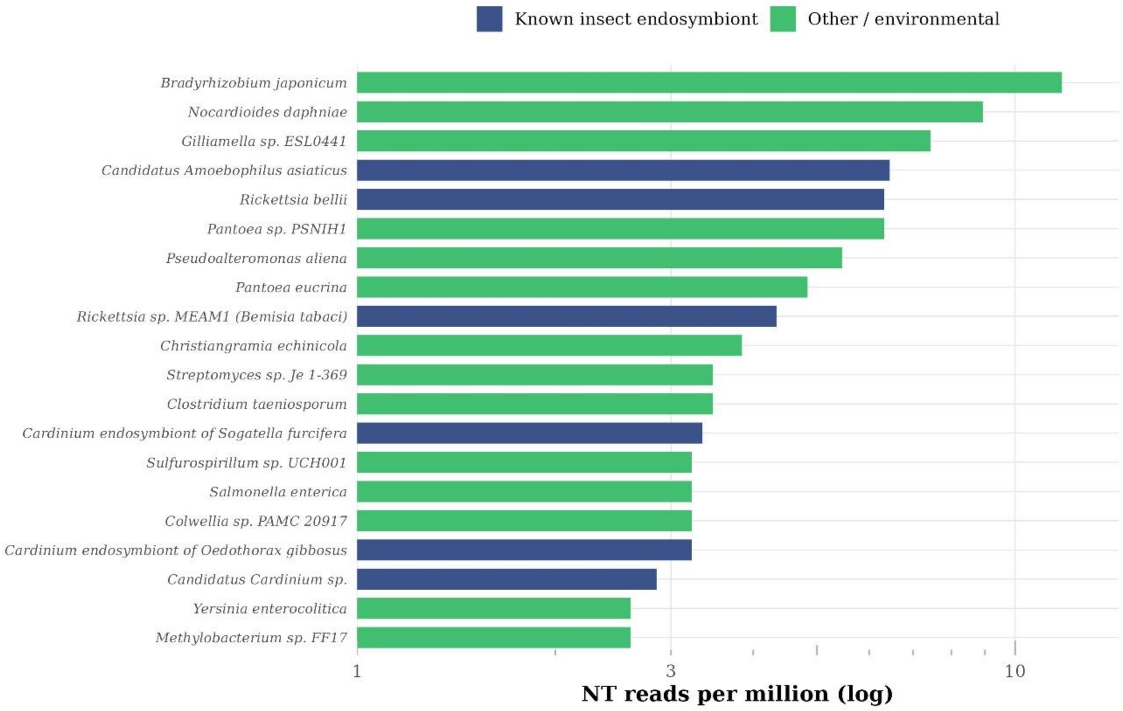


**Figure S2**. Bacterial microbiome with recognised insect endosymbionts highlighted. The most abundant bacterial taxa (NT reads per million, log); recognised insect-endosymbiont genera (Rickettsia, Cardinium, Arsenophonus, Wolbachia and the Cardinium-related Amoebophilus) are distinguished from other/environmental bacteria.
